# Synergistic anti-tumor activity, reduced pERK, and immuno-stimulatory cytokine profiles with 5-FU or ONC212 plus KRAS G12D inhibitor MRTX1133 in CRC and pancreatic cancer cells independent of G12D mutation

**DOI:** 10.1101/2023.09.13.557593

**Authors:** Vida Tajiknia, Maximilian Pinho-Schwermann, Praveen R. Srinivasan, Liz Hernandez Borrero, Leiqing Zhang, Kelsey E. Huntington, Wafik S. El-Deiry

## Abstract

KRAS mutations occur in ∼40-50% of mCRC and are associated with aggressive disease that is refractory to anti-EGFR therapies. Pancreatic cancer harbors ∼90% KRAS driver gene mutation frequency. Small molecules targeting KRAS G12C gained FDA approval for KRAS G12C-mutated NSCLC. ONC212, a fluorinated imipridone with nM anti-cancer activity has preclinical efficacy against pancreatic cancer and other malignancies. MRTX1133, identified as a noncovalent selective KRAS G12D inhibitor that suppresses G12D signaling by binding to the switch II pocket thereby inhibiting protein-protein interactions. We investigated cell viability, drug synergies, pERK suppression and cytokine, chemokine or growth factor alterations following treatment with 5-Fluorouracil (5-FU) or ONC212 plus MRTX1133 in 6 human CRC and 4 human pancreatic cancer cell lines. IC50 sensitivities ranged from 7 to 12 μM for 5-FU, 0.2-0.8 μM for ONC212, and >100 nM to >5,000 nM for MRTX1133 (G12D N=4: LS513 >100, HPAF-II >1,000, SNUC2B >5,000, PANC-1 >5,000). For non-G12D, the range of IC50 for MRTX1133 was >1,000 to >5,000 nM for CRC lines with G12V, G13D, or WT KRAS (N=7). Synergies between MRTX1133 plus 5-FU or ONC212 were observed regardless of KRAS G12D mutation with combination indices of <0.5 indicating strong synergy. Observed synergies were greater with MRTX1133 plus ONC212 compared to MRTX1133 plus 5-FU. pERK was suppressed with mutant but not wild-type KRAS at nM MRTX1133 doses. Immunostimulatory profiles included reduction in IL8/CXCL8, MICA, Angiopoietin 2, VEGF and TNF-alpha and increase in IL-18/IL-1F4 with MRTX treatments and combinations. Our studies reveal preclinical activity of MRTX1133 alone or synergies when combined with 5-FU or ONC212 against mCRC and pancreatic cancer cells regardless of KRAS G12D mutation. The results suggest that KRAS G12V and KRAS G13D should be further considered in clinical trials including combination therapies involving MRTX1133 and 5-FU or ONC212.

## Introduction

Colorectal cancer (CRC) is among the deadliest cancer and It is ranked the second in US cancer associated death.in 2023, in US, 153,020 people are projected to receive CRC diagnosis with 52550 total deaths attributed to this disease. Among these, 19,550 cases as well as 3750 fatalities are patients younger than 50.(1) The 5-year survival rate of Pancreatic cancer (PDAC) is around 12% in the US. With approximately 64,050 new cases in 2023 with projected 50,550 death in 2023, PDAC holds the fourth place in cancer related death. The incidence of pancreatic cancer has gone up around 0.2% year since 1990. Globally 466,003 people died from pancreatic cancer in 2020.(2–4)

KRAS is the most frequently mutated Oncogene in cancer.(5) Pancreatic, colorectal, lung adenocarcinomas and urogenital malignancies are the major cancers associated with KRAS mutation.(6) KRAS is a switch with intrinsic GTPase activity, it plays a major role in different cellular pathway such as proliferation and it modulates important signaling cascade such as MAPK pathway(7,8) when it is bound to GTP, KRAS becomes active, initiating the essential signaling pathways for proliferation and angiogenesis. KRAS Mutations keep the switch in a continuously ‘ON’ state. this broken switch now intensifies downstream signaling, resulting to formation of tumor.(8–11) targeting mutated KRAS has been one of the greatest challenges in cancer drug discovery. In fact, National Institutes of Health (NIH) has launched a special initiative to encourage focused effort on successfully targeting mutant KRAS, also known as ‘RAS initiative’(12)

The discovery of KRAS in 1982 is considered as the beginning of molecular oncology in human cancer research. (13,14) After almost 4 decades of considering KRAS as undruggable (15,16) in 2013 the ground-breaking work of Shokat and his team was published, (17) introducing molecules that bind to KRAS G12C covalently.(17,18) Different pharmaceutical companies followed this approach to introduce new KRAS G12C inhibitors.(19) The FDA granted accelerated approval to Sotorasib in May 2021 for the treatment of adult patients who have locally advanced or metastatic NSCLC with KRAS G12C mutation (20,21) and in December 2022 Adagrasib obtained its initial approval from FDA for treating adults with locally advanced or metastatic NSCLC harboring KRAS G12C mutation.(22) These two KRAS G12C inhibitors have in common is they both lock KRAS in an inactivate state.(23,24) A third agent, Divarasib (GDC-6036), that covalently targets KRAS G12C has recently been reported to have clinical activity across solid tumor types with durable clinical responses (DOI: 10.1056/NEJMoa2303810).

The problem with the G12C specific KRAS inhibitors has been toxicity with 11.6% of patients experiencing toxic effects in clinical trial CodeBreak100 (NCT03600883). (25) Another challenge has been resistance to KRAS G12C inhibitor, due to intrinsic or acquired resistance (26) from the CodeBreak 100 trial with median progression-free survival as low as 6.3 months.(25–27) In non-small cell lung cancer, data shows prevalence of different KRAS subtypes as G12C (40%), G12V (19%) and G12D (15%).(28) KRAS G12D is the most common driver mutation in PDAC and CRC.(29)

MRTX1133 was discovered through an extensive structure-based modification of KRAS G12C. This noncovalent potent inhibitor binds to both active and inactive forms of KRAS G12D and it is known to be a first-in-class KRAS G12D inhibitor.(30) The KRAS G12C Inhibitor efficacy relies on a stable covalent bond with mutant Cys12. (31,32) KRAS G12D protein lacks a reactive residue near the switch ll pocket (33) resulting a totally different mechanism of inhibition by MRTX1133 as compared to KRAS G12C inhibitors. We explored MRTX1133 synergies with current CRC and PDAC first-line treatments including 5-FU and whether there are effects on immune tumor microenvironment following MRTX1133 with or without combinatorial therapies. We explored combination therapies for targeting KRAS such as imipridone ONC212 and whether such synergies might be observed with non-G12D KRAS subtypes.

## Materials and Methods

### Tumor Cell Lines

Cells were cultured in specific medium (DMEM, EMEM, McCoy’s, RPMI) (high glucose without sodium pyruvate) supplemented with 10% heat-inactivated fetal bovine serum (FBS), and 1% penicillin/streptomycin. Human CRC and PDAC cell lines (BxPC-3, AsPC-1, and MIA PaCa-2) were purchased from ATCC. Cell lines were maintained according to ATCC guidelines.

### MRTX

MRTX1133 was purchased from Med Chem Express. The authenticity and purity of the chemical was verified by mass spectrometry as shown in the results.

### Cell Viability Assay

To determine IC50 values, 2 to 5 × 10^3^ tumor cells were plated in a 96-well plates. Cells were treated 24 hours later with DMSO or serial dilutions of MRTX1133. Cell viability was determined at 72 hours later using CellTiter-Glo (Promega, G7571). All reagents and plates were brought to room temperature before measuring viability. An equal volume of CellTiter-Glo was then added to each well. The cells were lysed, and plates were incubated for 10 minutes at room temperature. Luminescence values were recorded, and IC50 values were generated using GraphPad Prism (RRID:SCR_002798) version 9.1.2.

### Western Blot Analysis

Cells were washed with ice-cold PBS, and lysates were generated using RIPA buffer. Equal amounts of protein were run in reducing conditions on SDS-PAGE gels and then transferred to a PVDF membrane. Membranes were blocked at room temperature for 1 hour in a 5% Milk. After blocking, membranes were incubated in primary antibody diluted in 5%milk overnight at 4°C. After 3 washes in PBS-T, membranes were incubated at room temperature for 1 hour with fluorescence-conjugated secondary antibodies) diluted 5% milk. Primary antibodies for Western blot included anti–p-ERK (Cell Signaling Technology, 4370).

### Cytokine Assays

To assess secreted factors, tumor cell conditioned media was collected from cells treated with vehicle or MRTX1133 for 24 and 48 hours. Conditioned media was spun at 1,500 rpm to remove debris, and the supernatant was stored at −80°C. Conditioned media was analyzed using the custom panel of 60 cytokines (Luminex). Measurements were performed in duplicate.

### Organoid models

C57 BL/6 mice with truncated adenomatous polyposis coli (APC) floxed allele were crossed with heterozygous KRAS floxed outbred mice to generate an APCf/f KRAS+/f mouse colony. In another set of breeding, APC floxed mice were crossed with CDX2-Cre-ERT2 mice and selected for APCf/f CDX2-Cre-ERT2 after the second round of inbreeding. The final model of the disease was generated by the cross of the two parental colonies and viable APC f/f KRAS +/f CDX2-Cre-ERT2 (KPC: APC) were genotyped and characterized. The model animals are tamoxifen (TAM) inducible to generate tumors and overexpress KRAS G12D.

### Spheroid culture

A total of 70 μl complete media containing 1-2.4 × 10^3^ cells was seeded in each well in an ultra-low attachment 96-well plate (Corning® 96-well Clear Round Bottom Ultra-Low Attachment Microplate, NY, USA). Plates were centrifuged at 300 × g for 3 min and treated with the indicated compounds at different time points. Pictures were taken using ImageXpress.

Regarding the 3D structure of spheroid with necrotic core, quiescent zone and proliferating zone, effects on spheroid structure and proliferating zone was assessed in combination treatment in comparison to control in both KRAS G12V and KRAS G12D spheroids. Cell titerGlo assay was performed after transferring spheroid to 96 well black bottom plate.

### Statistical Analysis

GraphPad Prism (RRID:SCR_002798) version 9.1.2 was used for statistical analysis and graphical representation. For IC50 generation, concentrations were log-transformed, data were then normalized to control, and log(inhibitor) versus response (three parameters) test was used.

## Results

### MRTX1133 KRAS G12D inhibitor has inhibitory effects on non-G12D mutant subtypes but not on wild-type KRAS

Human PDAC and CRC cells with KRAS G12D, KRAS G13D, KRAS G12V and WT were cultured in 96-well plates overnight and treated with increasing concentrations of MRTX1133. At 72 hours, cell viability was determined using CTG luminescence. Western immunoblotting was used to investigate increasing concentrations of MRTX1133 on inhibition of pERK and this was shown at 24 hours. We unexpectedly discovered a potent effect of MRTX1133 on KRAS G12V as well as G13D, not just with regard to inhibition of cell viability but also towards reduction of pERK at low μM or nM doses of MRTX1133 (**Figure 1**). Cells with KRAS G12V showed sensitivity towards lowering pERK levels in the high nM or low μM doses of MRTX1133. Of note, KRAS wild-type (WT) cells showed resistance both to reduced cell viability or to reduction in pERK protein expression levels following MRTX1133 treatment.

**Figure 1.**
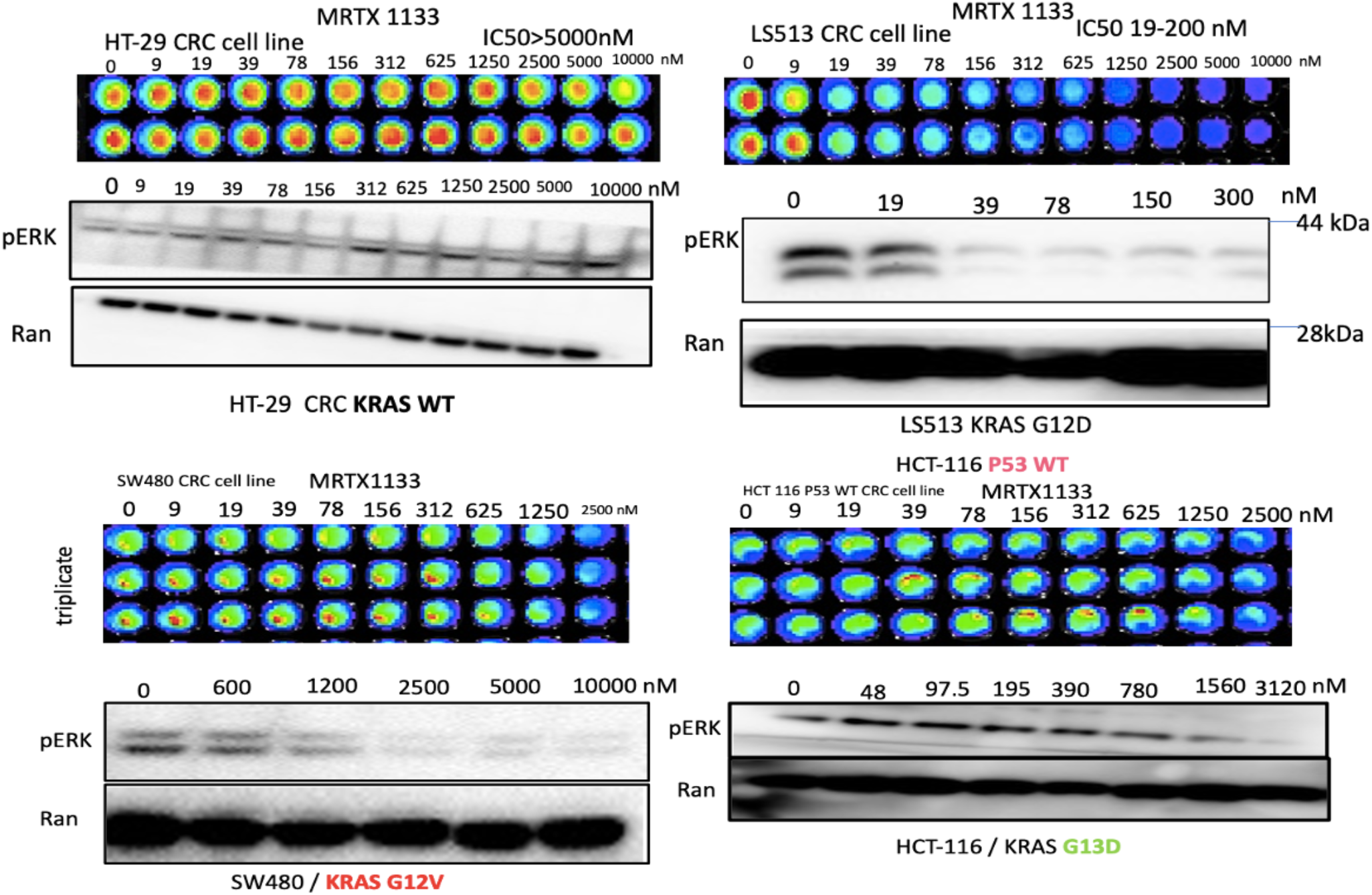
Sensitivity and pERK suppression in human CRC cell lines to MRTX as a function of KRAS mutation status. Cell viability assays with increasing MRTX1133 doses used to treat colorectal cancer cell lines with different KRAS mutations. The viability assays were performed using the CellTiterGlo assay. Western blots show expression of pERK and Ran with increasing doses of MRTX in the colorectal tumor cell lines as indicated. Ran was used as a loading control.

We further evaluated the sensitivity of human colorectal or pancreatic cancer cell lines with G12D, G12V or WT KRAS by determining IC50 after treatment with MRTX1133 (**Table 1**). We plated 2 to 5 × 10^3^ tumor cells in 96-well plates and treated the cells 24 hours later with DMSO or serial dilutions of MRTX1133. Cell viability was determined at 72 hours later using the CellTiter-Glo luminescence assay. We found that colorectal (LS513, SNU-C2B, HCY-116) or pancreatic (HPAF-II, PANC-1) tumor cell lines with KRAS G12D showed a range of IC50 values ranging from less than 50 nM to >1 μM while colorectal (SW480, SW620) or pancreatic CAPAN-1, CAPAN-1, CFPAC-1) tumor cell lines with KRAS G12V have a similar sensitivity near 1 μM. KRAS G13D tumor cells showed IC50 to MRTX1133 in the low μM range and also suppressed pERK in the nM range in preliminary experiments (**Figure 1**). By contrast to these mutant KRAS expressing human tumor cell lines, colorectal (HT-29) or pancreatic (BxPC-3) tumor cell lines with wild-type KRAS showed resistance to MRTX1133 and had much higher IC50 values >10 μM.

**Table 1.** Sensitivity (IC50, pERK suppression) of human CRC or PDAC cell lines independent of KRAS G12D driver mutation while KRAS WT tumor cells are resistant.

| Cell line | Kras status | pERK inhibition | MRTX1133 IC50 | Cell line | Kras status | pERK inhibition | MRTX1133 IC50 |
| --- | --- | --- | --- | --- | --- | --- | --- |
| LS513 | KRASG12D | 39nM | 39nM | CAPAN-1 | KRASG12V | 1.25 μM | 2.5-3.5 μM |
| SNU-C2B | KRASG12D | 612 nM | 1.25 μM | CAPAN-2 | KRASG12V | 612 μM | 1.25-2 μM |
| HCT-116 | KRASG13D | 780 nM | 2 μM | CFPAC-1 | KRASG12V | 1.25 μM | 2.5-4 μM |
| HPAF-II | KRASG12D | 1.2 μM | 1.77 μM | SW480 | KRASG12V | 612 nM | 2.79 μM |
| PANC-1 | KRASG12D | 2 μM | 2.5 μM | SW620 | KRASG12V | 1.2 μM | 3.6 μM |
| AsPC-1 | KRASG12D | 1.2 μM | 1.5-2.41 μM | HT-29<br>BxPC-3 | Wild type<br>Wild Type | - | >10 μM |

### Combination of MRTX1133 KRAS G12D inhibitor and 5-FU results in robust synergy with lower drug doses versus monotherapy

We examined synergies between MRTX1133 and the 5-FU chemotherapeutic agent used to treat colorectal or pancreatic cancer and observed synergy (**Figure 2**). Such combination has not been tested in any clinical trial and our preliminary data shows that the synergy is observed in both KRAS G12D (LS513 and SNUC2B cells), KRAS G12V Capan-2 tumor cells, or KRAS G13D HCT-116 cells (**Figure 2**)

**Figure 2.**
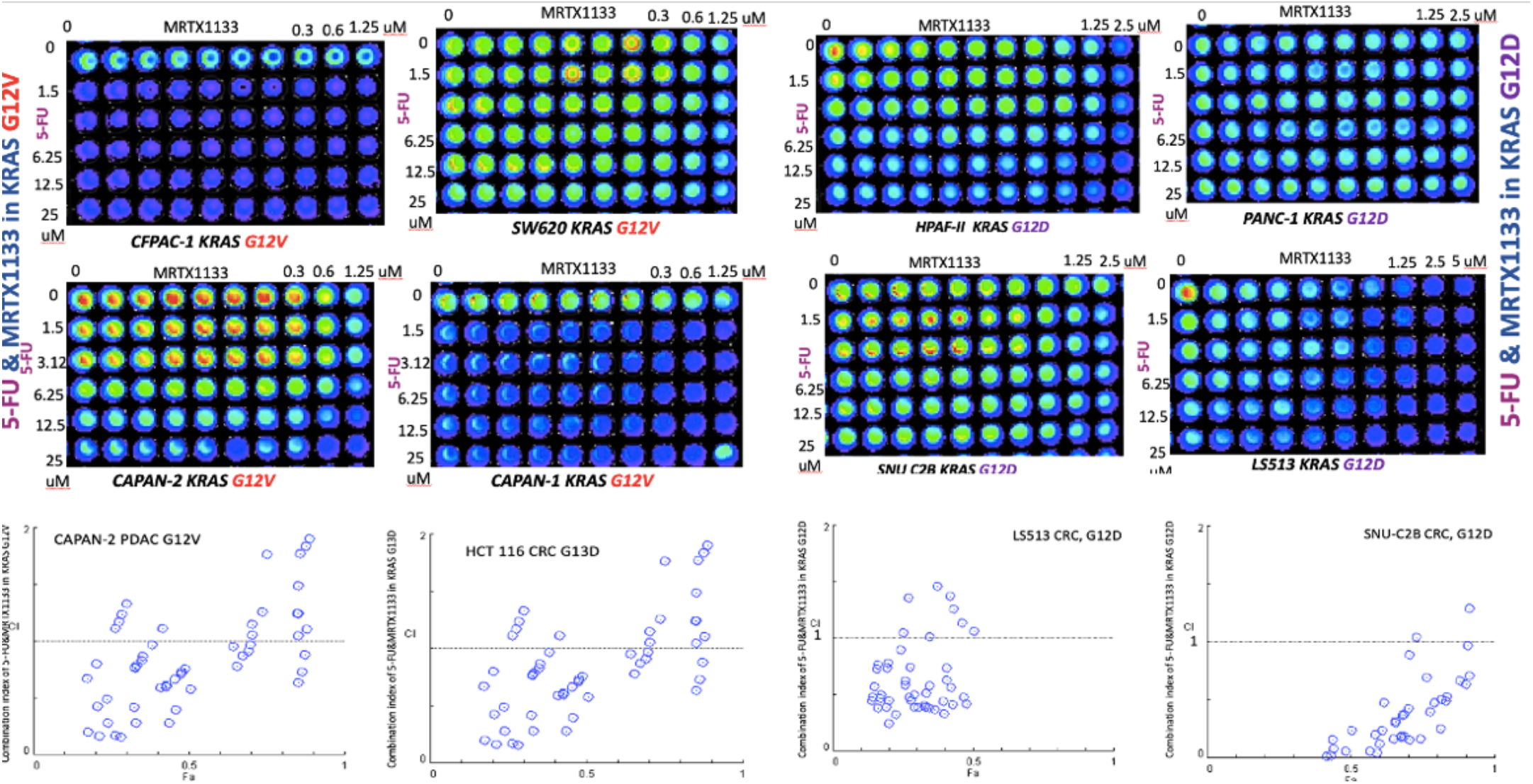
Synergy between 5-FU and MRTX1133 observed in KRAS G12V, KRAS G13D and KRAS G12D but not in KRAS WT CRC or PDAC cell lines, independent of KRAS mutation subtype. Strong synergy was observed between 5-FU and MRTX1133 in both KRAS G12V and KRAS G12D cell lines. Combination index ranges from 0.1-0.8 A: CI in LS513 (KRAS G12D), B: CI in SNUC2B (KRAS G12D), C: CI in Capan-2 (KRAS G12V), D: CI in HCT-116 (KRAS G13D) E&F, 72 hours cell viability assay used in cells with KRAS G12V & KRAS G12 D to determine the synergy. CI < 1 reflects drug synergy.

### Combination of ONC212 and MRTX1133 shows synergy and suppression of pERK regardless of KRAS G12D mutation status in CRC or PDAC

We examined synergies between MRTX1133 and ONC212 (**Figure 3**). We plated 2 to 5 × 10^3^ tumor cells in 96-well plates and treated the cells at 24 hours later with DMSO or serial dilutions of MRTX1133 and ONC212. Cell viability was determined at 72 hours later using the CellTiter-Glo assay. The results show synergistic effects of the MRTX1133 plus ONC212 combination against the CRC and PDAC cell lines. Synergistic inhibition of pERK was observed in SW480 KRAS G12V, CAPAN2 KRAS G12V as well as the LS513 KRAS G12D cell line (**Figure 4**).

**Figure 3.**
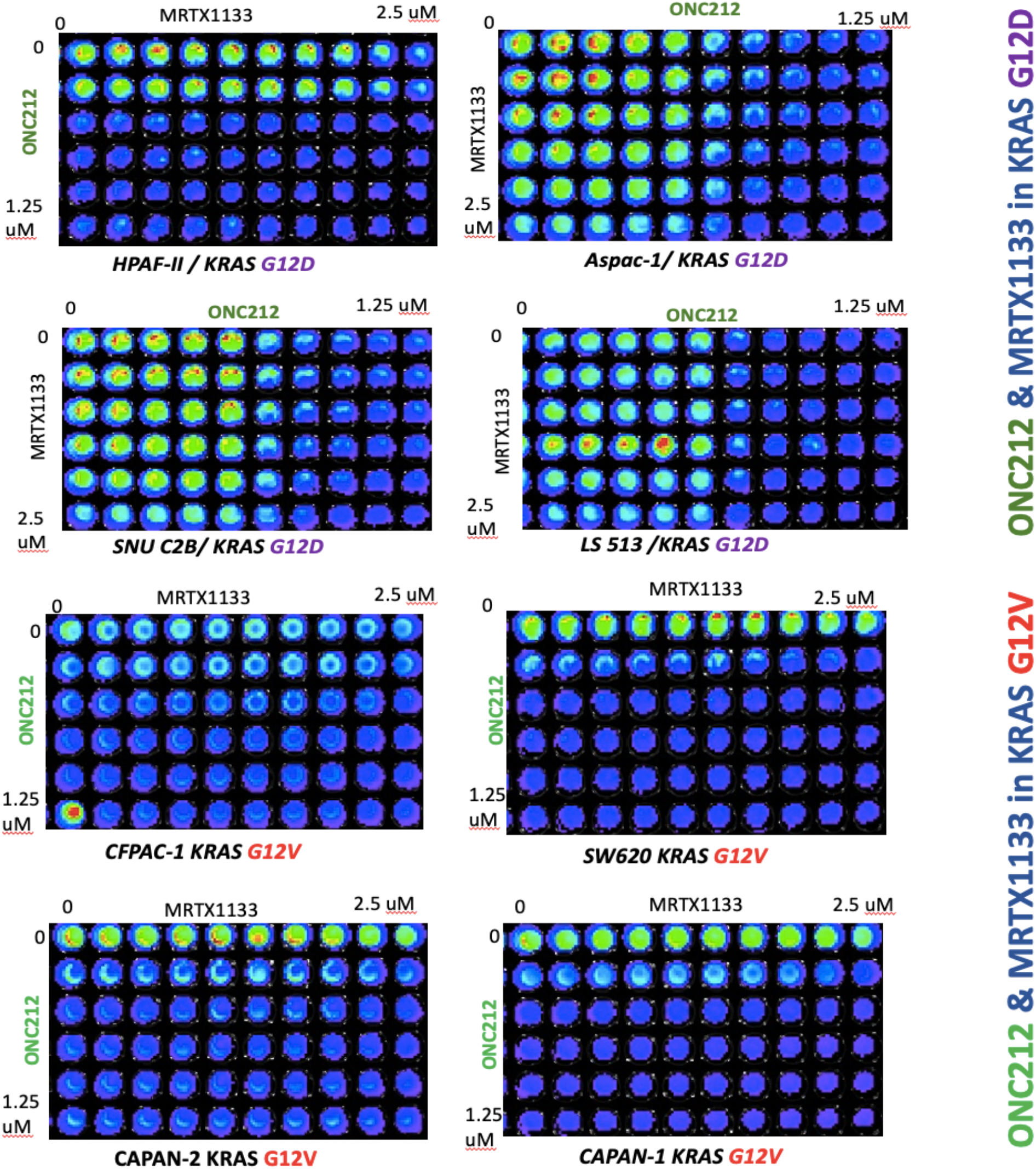
Synergistic effects of the MRTX1133 plus ONC212 combination against the CRC and PDAC cell lines.

**Figure 4.**
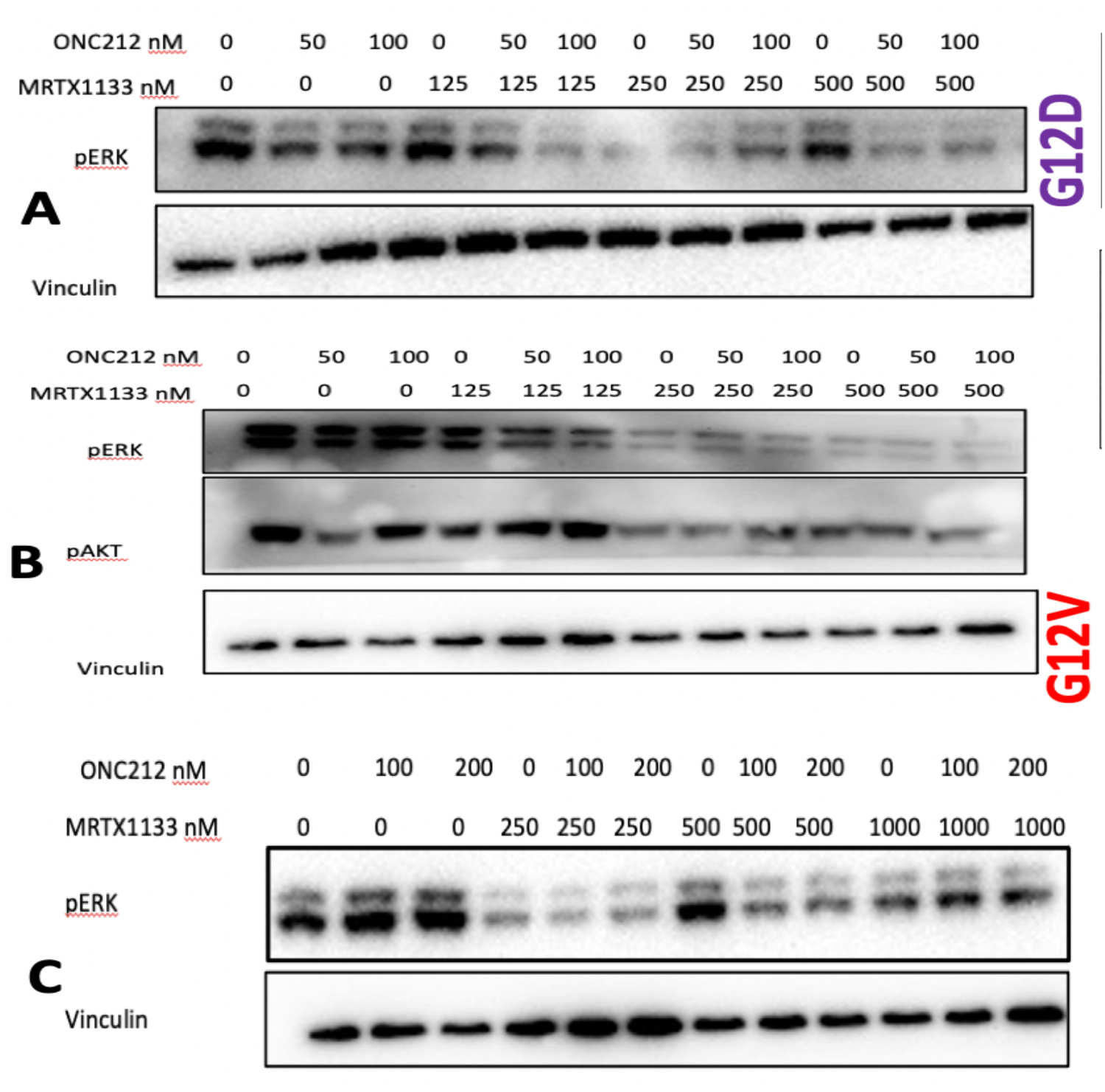
Synergistic inhibition of pERK with MRTX1133 and ONC212 in KRAS G12D and G12V cell lines. (A) G12D LS513 cell line, (B) Capan-2 KRAS G12V (C) KRAS G12V SW480 cell line, Western blot, 24 hours.

### MRTX1133 induces tumor suppressive immune microenvironment cytokine profiles in KRAS G12D as well as KRAS G12V sub-types

We examined bulk-tumor cell cytokine profiles after treatment of tumor cells harboring different KRAS mutation sub-types with 5-FU and MRTX1133 monotherapy or combinations (**Figure 5**). To assess secreted factors, tumor cell-conditioned media was collected from cells treated with vehicle or MRTX1133 for 24 and 48 hours. Measurements were performed in duplicate. We observed an increase in IL-18 after combination therapy regardless of KRAS mutation status, and a decrease in IL-8, MICA, VEGF, Angipoietin2 and TGF-Alpha at the 48-hour time point (**Figure 5**).

**Figure 5.**
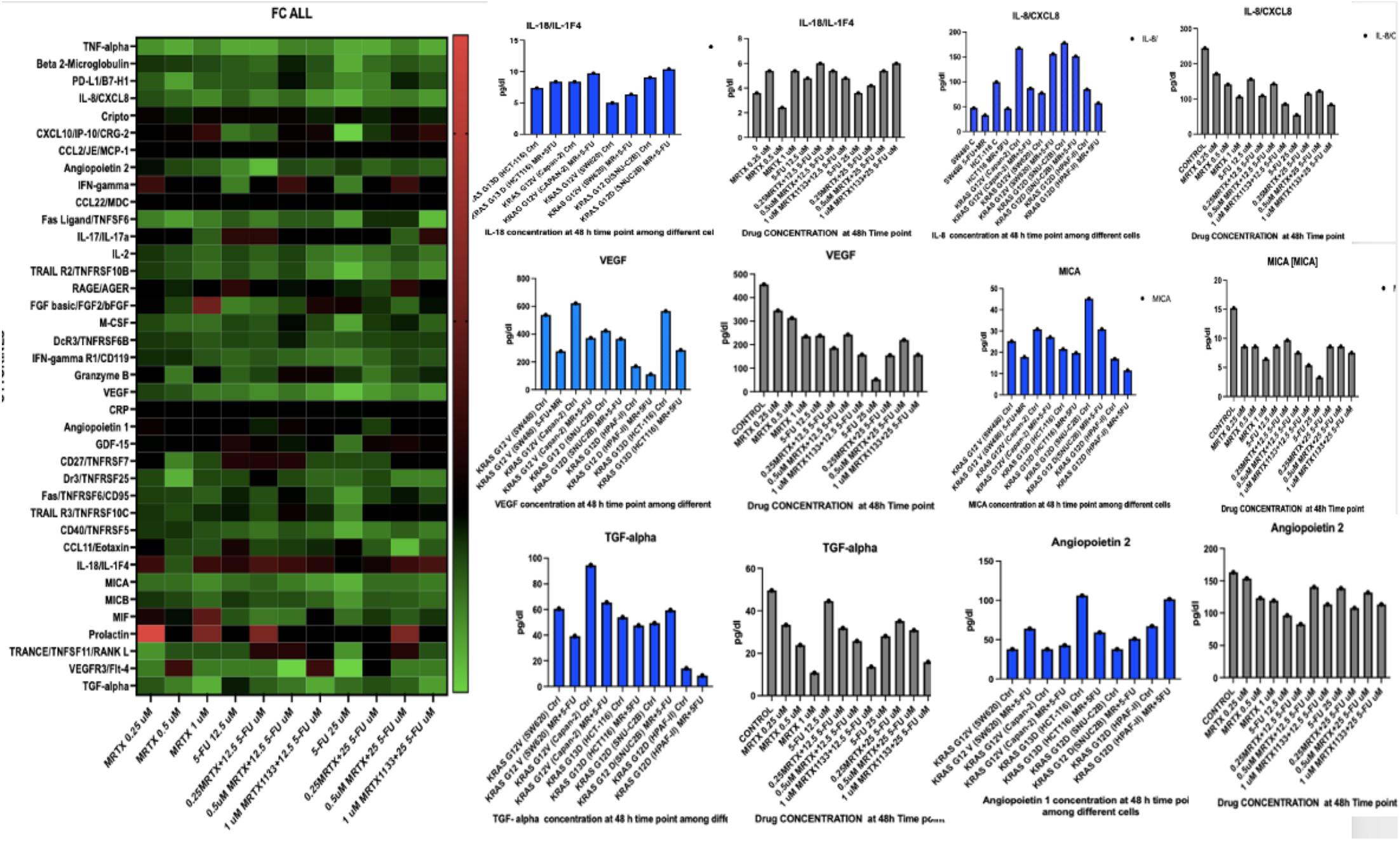
Cytokine Analysis in KRAS G12D vs KRAS G13D and G12V treated with 5-FU and MRTX1133, Effects are independent of KRAS subtypes with no effect on wild-type (WT). Increase in IL-18 after combination therapy was observed regardless of KRAS mutation status, also a decrease in IL-8, MICA, VEGF, Angiopoietin 2 and TGF-Alpha was found at 48-hour time point.

### Synergy observed in transgenic mouse organoids and spheroid cultures harboring both KRAS G12V and G12D mutations

We examined 3D spheroid structure and observed a necrotic core, quiescent zone and proliferating zone destruction of spheroid structure and smaller proliferating zone in combination treatment versus control in both KRAS G12V and KRAS G12D spheroids (**Figure 6**). CellTiterGlo assay revealed strong synergy in reduction of cell viability with combination of MRTX1133 and ONC212.

**Figure 6.**
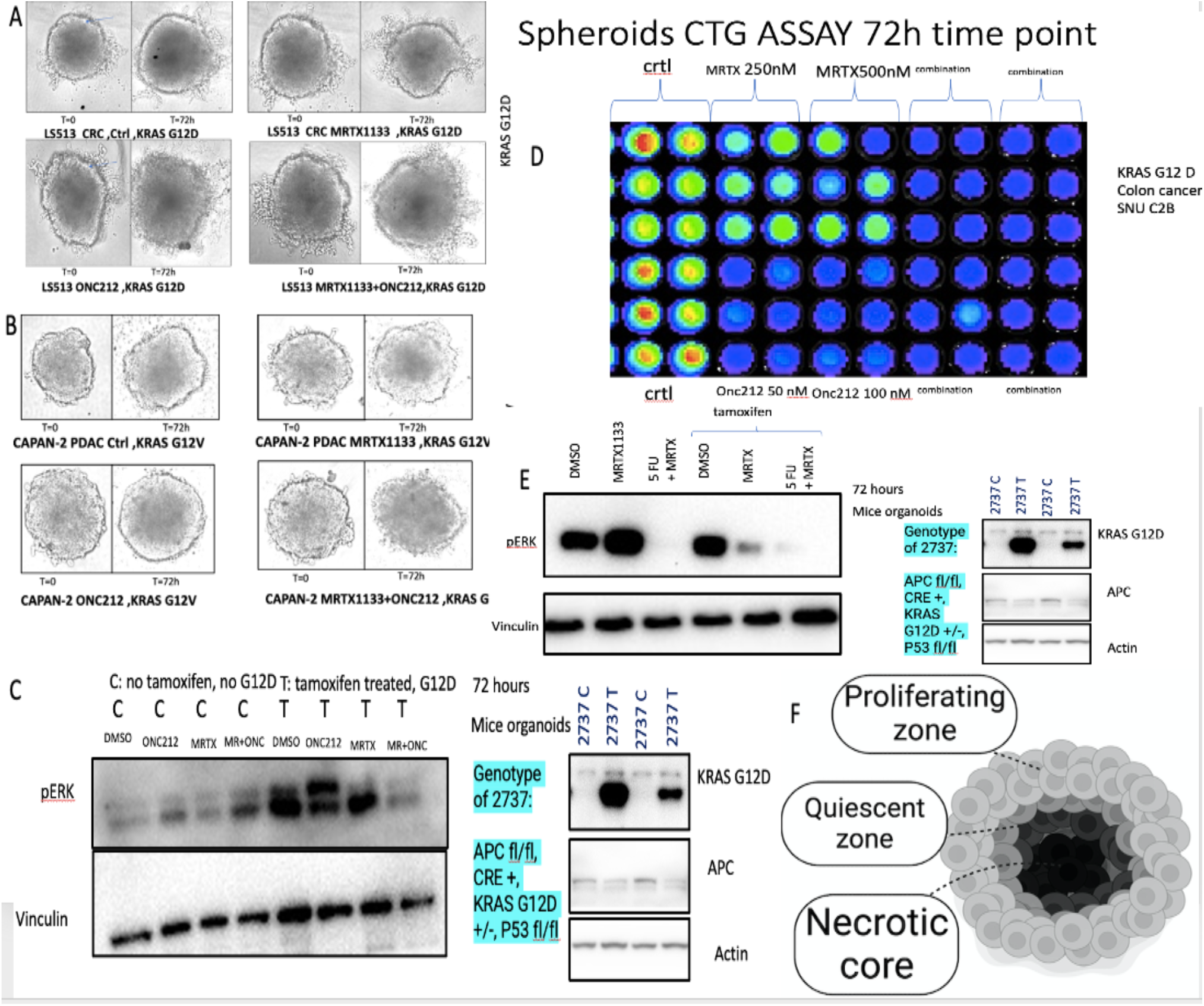
MRTX1133 treatment of 3D culture and transgenic mouse organoid models. (A, B) 3D spheroid structure and synergistic drug effects on spheroid models with G12D and G12V with MRTX1133 and ONC212. (C) Synergistic inhibition of pERK in treated (T) versus control (C) tumor cells. (D) Synergistic suppression of pERK after 5-FU and MRTX1133 treatment of transgenic organoids with tamoxifen inducible knockout of APC, knock-in of KRAS G12D, and p53+/-(left) or tamoxifen-inducible knockout of APC, knock-in of KRAS G12D, and p53+/-(right) at 72 hours. (E) 72h CellTiterGlo loss of cell viability of KRAS G12D spheroids with combinations of MRTX1133 and ONC212 as an example.

Tamoxifen induced KRAS G12D knock-in model was used to investigate effect of MRTX1133 and combination therapy on organoids with inducible oncogenic alleles for APC, p53, and KRAS. APC floxed mice were crossed with CDX2-Cre-ER^T2^ mice and selected for APC^f/f^ CDX2-Cre-ER^T2^ after the second round of inbreeding. The final model of the disease was generated by the cross of the two parental colonies and viable APC ^f/f^ KRAS ^+/f^ CDX2-Cre-ER^T2^ (KPC: APC) were genotyped and characterized. The model animals were tamoxifen (TAM) induced to generate tumors. We chose this model is an excellent preclinical platform to molecularly characterize the KRAS mutated colorectal tumors and discern appropriate therapeutic strategies to improve disease outcomes.

As shown in **Figure 6** synergistic inhibition of pERK in transgenic organoids treated with Tamoxifen (T) versus control (C) tumor cells was observed. Synergistic suppression of pERK after 5-FU and MRTX-1133 treatment of transgenic organoids with tamoxifen inducible knockout of APC, knock-in of KRAS G12D, and p53+/-or tamoxifen-inducible knockout of APC, knock-in of KRAS G12D, and p53+/-at 72 hours was observed.

### MRTX1133 Combination with ONC212 enhances NK-cell mediated death in KRAS G12D expressing tumor cells

To evaluate immune-mediated killing of tumor cells expressing mutated KRAS, we performed immune co-culture incubation assays using NK cells and LS513 KRAS G12D cells (**Figure 7**). Tumor cells were dyed as blue, immune cells as green, and ethidium homodimer was added to show cell death in red. The average number of dead cells was measured. The highest death belonged to combination therapy with NK cells.

**Figure 7.**
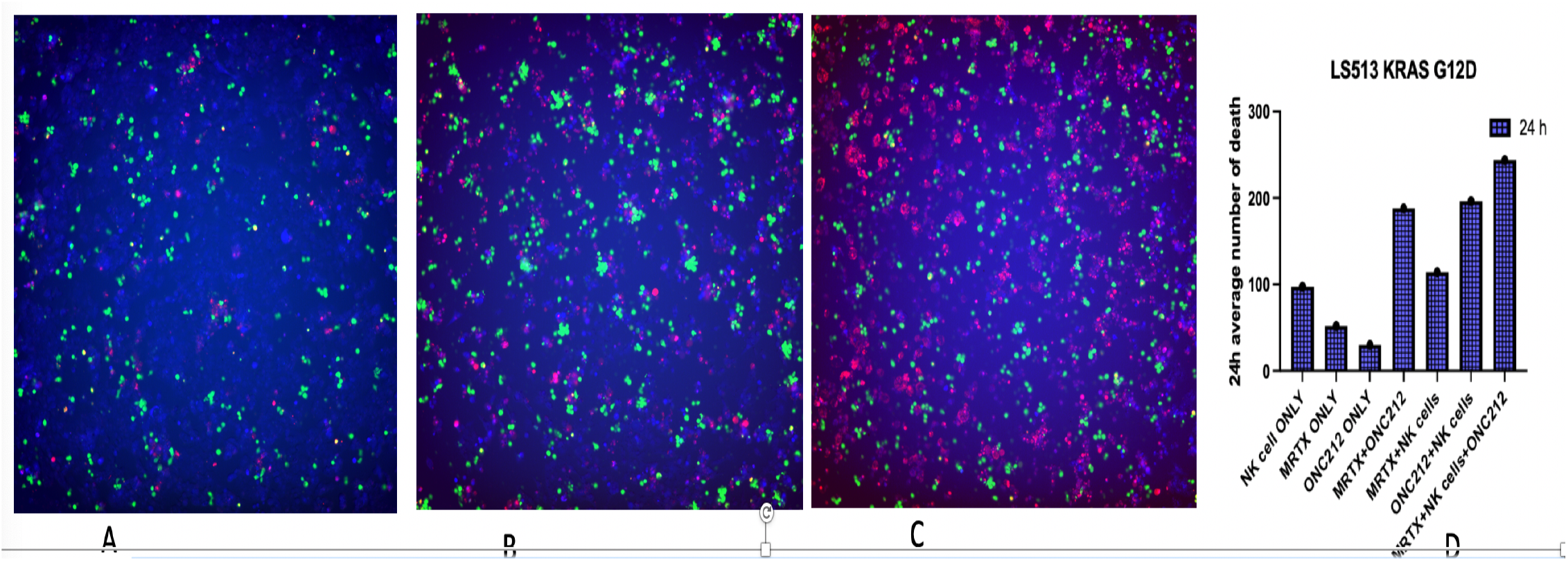
Immune Co-Culture with NK cells at 24 h showed increase in cell death with MRTX1133, ONC212 and synergy in combination with NK cells compared to without immune cells. In KRAS G12D cell line the result of immune co-culture showed enhanced NK mediated cell death which is more significant in combination

### MRTX1133 verification

Since we obtained the drug commercially through an available vendor, we assessed drug authenticity and purity. We used Mass spectrometry to investigate drug authenticity (**Figure 8**). Data showed the drug is almost 100% the one that published data of MRTX1133 is claimed to be.

**Figure 8.**
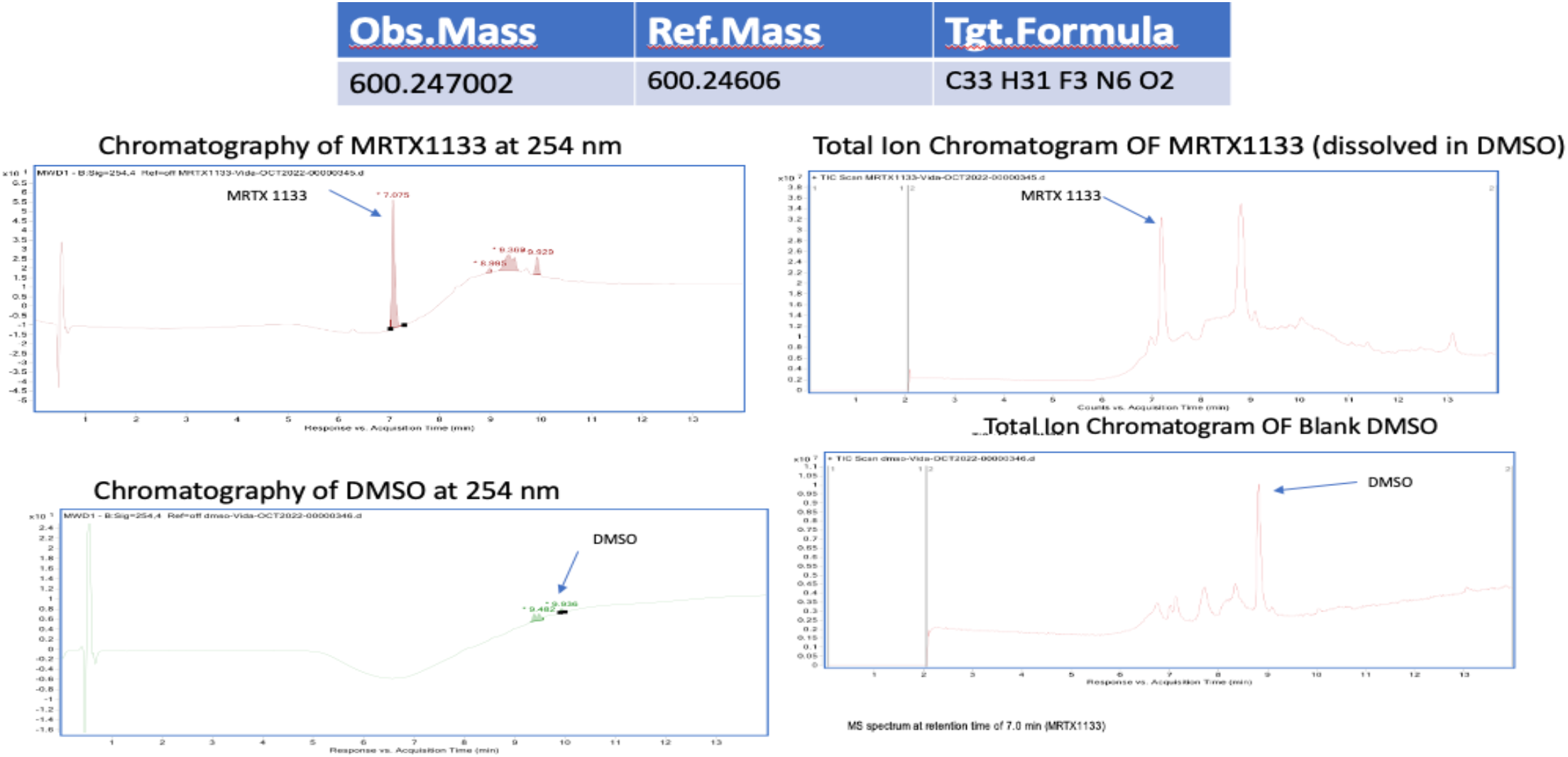
The molecular weights measured for the commercially available chemical highly match with theoretical MW of MRTX1133. Evidence to determine the presence and authenticity of molecule mass of MRTX1133 by mass spectrometry. Compare the TIC and UV chromatography of MRTX1133 and blank DMSO, no obvious impurities observed.

## Discussion

Targeting oncogenic mutant KRAS proteins in tumor cells has been shown to be one of the greatest challenges in cancer research and oncology. With the recent development and clinical success of KRAS G12C inhibitors in lung cancer, KRAS G12D inhibitor MRTX1133 was identified. Our results suggest that non-G12D KRAS mutant subtypes might benefit from the use of MRTX1133 including the common KRAS G12V mutation. We found synergy between standard of care 5-FU and the MRTX1133 KRAS G12D inhibitor as well as a synergistic effect of ONC212 and MRTX1133 as novel combination for targeting KRAS as an objective of the RAS initiative. The key finding here is the effects of MRTX1133 and synergistic combinations extended beyond KRAS G12D and were not limited to growth inhibition or downregulation of pERK, but also to meaningful impact on immune response. MTRX stimulated killing of tumor cells by NK cells and promoted changes in cytokine profiles to a favorable immune stimulatory microenvironment.

Small molecules have their challenges with resistance and toxicity. Our hypothesis was if MRTX1133 can be added to standard-of-care therapeutics it may be possible to achieve synergy and efficacy at lower and less toxic doses of drugs used in combination. So we expected a smaller dose of both drugs such as MRTX1133 plus 5-FU or MRTX1133 plus ONC212 will be required for the cytotoxic response. Additionally, we suspect that use of a synergistic combination will reduce the chances for development of resistance to therapy at an early stage of treatment and this may translate to a better clinical outcome for patients.

It worth noting that different cells with the same KRAS G12D mutation demonstrated different responses to MRTX1133 (different IC50s and different immune microenvironment changes). It is likely that other genetic or epigenetic alterations may explain this. We do not know but suspect that co-mutation with p53 might affect sensitivity to MRTX1133 especially when combined with agents such as 5-FU that activate p53 signaling. We suspect that other mutations including those that may be different between CRC and PDAC may impact response to therapy as well.

To this date published data regarding MRTX1133 showed efficacy against only KRAS G12D and effect on T cells. (30, 34–39)

Since MRTX1133 binds non-covalently to KRAS G12D in both the active and inactive form, the key to maximizing effectiveness of this small molecule in KRAS G12V cells must be a non-covalent interaction.

Our results are limited by lack of *in vivo* data. Future studies are required to investigate the *in vivo* correlates, but we did examine organoids and 3-D cultures in the present study. Currently the NCT05737706 phase 1 clinical trial ‘Study of MRTX1133 In Patients With Advanced Solid Tumors Harboring a KRAS G12D Mutation’ does not include patients whose tumors harbor KRAS G12V or other KRAS non-G12D mutations. The importance of our finding is to consider treatment of tumor with other KRAS mutations (no information currently on NRAS) and by introducing combination therapy as a rational strategy against toxicity and resistance. The immune-stimulatory effects observed in our study require further analysis and experimentation to determine the utility and predictive value as far as response to clinical therapy. Our results suggest combinations of MRTX1133 with 5-FU or ONC212 for further study in preclinical models as well as in the clinic.

## Acknowledgements

This work was presented in part at the April, 2023 American Association for Cancer Research (AACR) meeting in Orlando, and the 12th AACR-JCA Joint Conference on Breakthroughs in Cancer Research: Translating Knowledge into Practice, December 10-14, 2022, Maui, Hawaii. The work was supported by a Warren Alpert Foundation grant (to W.S.E-D.), an NIH grant to W.S.E-D. (CA173453), a grant from Chimerix, Inc. to W.S.E-D. and the Mencoff Family Professorship at Brown University (W.S.E-D.). W.S.E-D. is an American Cancer Society Research Professor.

